# Chronic oral fentanyl consumption produces dynamic behavioral adaptations during abstinence in female mice

**DOI:** 10.64898/2026.07.27.741002

**Authors:** Mengfan Xia, Brianna E. George, Maximilian Caza, Jobe L. Ritchie, Thomas L. Kash

## Abstract

**Rationale:** The opioid epidemic continues to be driven by synthetic opioids, particularly fentanyl, yet the long-term behavioral manifestations of withdrawal remain poorly characterized. Women exhibit unique vulnerabilities to opioid use disorder, including greater susceptibility to withdrawal-related symptoms and relapse, suggesting that defining the behavioral adaptations that emerge during prolonged fentanyl abstinence may identify mechanisms underlying female-specific relapse vulnerability. Thus, we used an oral fentanyl drinking model to longitudinally examine nociceptive, affective, and exploratory behaviors across chronic exposure and abstinence in female mice.

**Methods:** Adult female C57BL/6J mice underwent a modified 5-week Drinking-in-the-Dark paradigm with 4-hour daily access to fentanyl (30 µg/mL) or water. Somatic withdrawal was assessed following naloxone-precipitated withdrawal. Thermal and mechanical nociception, sucrose preference, exploratory behavior, acoustic startle, nest building, and locomotor activity were evaluated during fentanyl exposure and throughout one month of abstinence.

**Results:** Female mice exhibited robust naloxone-precipitated somatic withdrawal, confirming physiological effects of opioid drinking. Chronic fentanyl exposure produced transient thermal hyperalgesia during weeks 2–4 of drinking that resolved by the final week, but re-emerged after 30 days of abstinence. After 30 days of abstinence, we also found increased mechanical sensitivity. During early abstinence, fentanyl-exposed mice exhibited increased sucrose consumption and greater exploration of anxiogenic environments without alterations in general locomotion. Increased exploratory behavior persisted into prolonged abstinence and was partially rescued in mice who previously received naloxone. Additionally, fDID mice exhibited impaired nesting behavior, while prior naloxone-precipitated withdrawal improved nest-building performance.

**Conclusions:** We demonstrate that chronic opioid exposure, precipitated withdrawal trials, and duration of abstinence interact to shape protracted withdrawal phenotypes in female mice that extend well beyond drug exposure. These results provide insight into persistent withdrawal symptoms that may shape relapse vulnerability using a novel translationally relevant framework.

## Introduction

The ongoing opioid crisis remains a major public health concern with synthetic opioids, particularly fentanyl, driving increased overdose deaths over the last decade^1^. Although opioid use disorder (OUD) affects both sexes, women exhibit unique vulnerabilities throughout the addiction cycle, including more rapid progression from initial use to dependence (telescoping^2^), higher drug craving^3^, and greater susceptibility to relapse^4^. Moreover, withdrawal symptoms are among the strongest predictors of future opioid relapse in females, highlighting the importance of understanding the behavioral consequences of opioid withdrawal in women^4^. While acute withdrawal symptoms are well characterized, considerably less is known about the persistence of affective and motivational disturbances during protracted fentanyl abstinence. Deciphering how behavioral manifestations develop across both acute and prolonged fentanyl withdrawal may serve to inform the development of sex-specific prevention strategies and pharmacotherapeutic interventions for OUD.

Prior evidence has shown that females are more likely to ingest opioids orally than males^5^. Further, rates of alternative methods of administration such as inhalation and ingestion have increased since 2015^6^ and the availability of fentanyl in tablet or pill form has also rapidly expanded^7–9^. While most preclinical studies have relied on experimenter-administered opioids or intravenous self-administration paradigms^10,11^, these models may not capture the patterns of volitional oral fentanyl use that have become increasingly common with the widespread availability of counterfeit fentanyl-containing pills^12^. More recently, home-cage oral fentanyl drinking paradigms have been developed using the established drinking-in-the-dark (fDiD) approach in which mice consume fentanyl over several weeks, producing pharmacologically relevant intake and robust physical dependence^13,14^. This model recapitulates several clinically relevant features of OUD, including volitional drug consumption, naloxone-precipitated withdrawal, disrupted sleep, alterations in reward-related behaviors, impaired fear extinction, and anxiety-like phenotypes during abstinence^13^. Notably, female mice consumed greater amounts of fentanyl than males at higher fentanyl concentrations in this paradigm, further supporting its utility for investigating mechanisms that may underlie sex-dependent vulnerability to OUD. However, the temporal profile of the development of withdrawal-associated behavioral disturbances in females remains incompletely characterized.

Negative affective states that emerge during opioid abstinence are known drivers of continued use and relapse, consistent with theories of negative reinforcement in addiction^15^. Acute opioid withdrawal can be characterized by somatic and affective symptoms that typically resolve within 7 days; protracted withdrawal is characterized by mostly psychological and affective symptoms that occur well beyond physical dependence to the substance^15,16^. Reviews of preclinical opioid abstinence models have highlighted enduring disruptions in anxiety-like behavior, stress responsivity, pain sensitivity, sleep, cognition, and reward processing that can persist for weeks or months following opioid discontinuation, yet these findings are derived predominantly from studies in males and vary largely across opioid used, route of administration, and duration of abstience^10,16–18^. Furthermore, relatively few studies have systematically tracked behavioral changes longitudinally throughout prolonged abstinence, making it difficult to distinguish transient, early withdrawal effects from persistent adaptations that may contribute to relapse vulnerability.

Therefore, the present study employed a modified home-cage fDiD model in female mice to characterize the emergence and persistence of sensitivity to noxious and non-noxious aversive stimuli, anxiety-like behaviors, and affective disruption across one month of prolonged abstinence. By defining the temporal profile of these behavioral adaptations, this work aims to improve our understanding of the mechanisms that contribute to relapse vulnerability in females and provide a translational framework for developing sex-specific therapeutic strategies for OUD.

## Methods

### Subjects

Adult female (n = 40) C57Bl/6J mice (The Jackson Laboratory, Bar Harbor, ME) were singly housed prior to the start of experiments. Home cages were ventilated in a colony room on a 12:12-hour reverse light-dark cycle (lights off at 0700). All animals had food and water *ad libitum* for the duration of the study. After the completion of the study, all animals were euthanized according to IACUC protocols. All procedures were approved by the Institutional Animal Care and Use Committee of the University of North Carolina at Chapel Hill and performed in accordance with the National Institute of Health’s guide for the care and use of laboratory animals.

### Drugs

Fentanyl citrate was purchased from Spectrum Pharmacy Products (New Brunswick, NJ, USA) and dissolved in tap water for drinking studies at dose of 30µg/mL. For precipitated withdrawal studies, naloxone hydrochloride was purchase from Sigma-Aldrich (St. Louis, MO, USA), dissolved in 0.9% physiological saline, and injected intraperitoneally in mice at a dose of 1mg/kg.

### Fentanyl Drinking in the Dark (fDiD)

At the start of the paradigm, mice were provided access to a single bottle containing fentanyl (30 µg/mL in tap water) in the homecage three hours into the start of the dark cycle (1000). Water control mice received a bottle of tap water alone. After four hours, the bottles were removed and replaced with drinking water until the next day. fDiD drinking sessions were run for 5 consecutive days with 2 days off for a total of 5 weeks. All bottles were weighed at the beginning and end of each drinking session to calculate the amount consumed. To account for dripping, one bottle of drinking water was placed on an empty cage and drinking and this value was subtracted from all daily drinking amounts. Mice were weighed weekly after the final fDiD session for that week.

### Precipitated Withdrawal Opioid Behaviors

Opioid withdrawal behaviors were quantified as previously described^13,19,20^. Following the final drinking session during the 4^th^ week of fDiD, mice were injected with either 1 mg/kg naloxone or saline control (N=10 per group) and placed in an open arena. Withdrawal behaviors were observed for 10 mins. The withdrawal behaviors assessed included escape jumps, paw tremors, jaw tremors, abnormal postures, wet dog shakes, grooming behaviors, and fecal boli. For normalization, data were converted to a z-score for each behavior and then averaged to generate a global somatic withdrawal score for each subject.

### Hargreaves

We used the Hargreaves test to assess thermal nociceptive sensitivity. Mice were habituated to the testing room for 30 minutes, then put in clear Plexiglas boxes on a raised glass surface and acclimated to the behavioral equipment for a minimum of 30 minutes. After that, each hind paw was subjected to a series of heat trials aimed at the mid-plantar surface, separated by a 10-minute interval. Radiant heat exposures were applied to the left and right paws sequentially during the trials. Four trials were conducted and averaged for each mouse. Beam intensity was set at 22 on the IITC Plantar Analgesia Meter (IITC Life Science, Woodland Hills, CA), which produced baseline paw withdrawal latencies around 10 seconds in this cohort of mice. To avoid tissue injury, a 25-second cutoff limit was applied.

### Hot Plate

Mice were placed in a custom-made cylindrical Plexiglas container on an IITC Hot Plate Analgesia Meter (IITC Life Science, Woodland Hills, CA), set on a black anodized aluminum plate measuring 28 cm x 28 cm. Mice were exposed to 55°C hot plate for 45 seconds to induce sensory-discriminative and affective-motivational behaviors linked to pain. This extended 45 s test allows for readout of behaviors beyond latency to the first paw withdrawal^21^. Computer-integrated Logitech cameras were used to record each session, allowing for blinded manual scoring of pain-related behaviors. Videos were scored for: paw withdrawal (rapid flicking of the limb), paw attending (licking of the limb), and paw guarding (intentional lifting of the limb for protection), and escape jumps.

### Von Frey Test

To evaluate mechanical reflexive sensitivity, we used the Von Frey test as previously described^22^. The mice were placed in clear Plexiglas boxes on a raised metal wire platform (90 × 20 × 30 cm) and habituated to the behavioral testing apparatus for at least 60 minutes. Using the simplified up-down method (SUDO)^23^, a set of 10 nylon monofilaments with forces ranging from 0.008 g to 2 g were applied perpendicularly to the hind paws. The test starts with a mid-range force of 0.16 g. The filament was applied to the mid-plantar surface of the hind paw for 10 trials. Each mouse received 10 trials with each filament on one paw, which was randomly assigned before the tests. Depending on the number of paw withdrawals (positive responses) when the filament bent, the procedure was then repeated with either increasing or decreasing forces. Each filament was tested for ten trials. The minimum gap time was 15 minutes between each filament. The minimum force (grams) at which a filament resulted in a withdrawal reflex in more than half (>=5) of the 10 trials was calculated as the 50% withdrawal threshold score.

### Open Field Test

The open field test (OFT) was conducted in a 51 cm x 51 cm opaque white Plexiglas box with an open top. The open field was illuminated from above by LED strips (100 lux at center). Mice were placed in the corner with their heads facing the center. Mice were tracked using Noldus EthoVision (Noldus Information Technologies, Wageningen, Netherlands). A 70% ethanol solution was used to clean the open field between subjects.

### Sucrose Preference Test

Mice were given access to water and 1% w/v sucrose in water for 24 hours in their home cage. Sucrose and water bottles were weighed before and after the test. The bottles were replaced with drinking water after testing. Sucrose consumption was normalized to body weight and a preference for sucrose over water was calculated.

### Elevated Plus Maze

Mice were placed in the center of the apparatus with their heads facing the opening of the closed arm and allowed to freely explore for 10 minutes. The maze was illuminated from above by two dim LED strips (open arm lux of 25) mounted to the ceiling. Sessions were video recorded and mice were tracked using EthoVision behavior tracking software (Noldus, Netherlands). A 70% ethanol solution was used to clean the maze between subjects.

### Acoustic Startle

The acoustic startle response was assessed using the SR-LAB Startle Response System (San Diego Instruments, San Diego, CA, USA). Mice were placed individually into a clear acrylic cylindrical restraint chamber positioned within a sound-attenuating chamber and allowed to acclimate for 5 min with continuous background white noise (65 dB). Following habituation, mice were presented with acoustic startle stimuli of 90, 105, and 120 dB (40 ms duration) in a pseudorandom order with variable intertrial intervals of 30–50 s. Each stimulus intensity was presented 10 times. The startle response was recorded for 200 ms following stimulus onset, and peak startle amplitude was defined as the maximum response occurring within the first 100 ms after stimulus presentation. Mean startle amplitude for each stimulus intensity was calculated by averaging responses across the 10 trials.

### Elevated Zero Maze

The elevated zero maze (EZM) assay was conducted on a circle shaped apparatus measuring 62 cm in outlining diameter, divided into four equal quadrants. Two opposite quadrants were “open”; the remaining two “closed” quadrants were surrounded by dark, opaque walls. Additional illumination for open areas was provided by LED strips lighting from the center of the apparatus plane. Each mouse was placed at the intersection of open and closed areas at the beginning of each trial, and an overhead camera recorded activity for 10 min. Time spent and distance traveled, moving velocity in each area was analyzed with EthoVision software (Noldus, Netherlands). A 70% ethanol solution was used to clean the maze between subjects.

### Nestlet Shredding and Nest Building Assay

Both assays were performed according to previously published protocols^24,25^. Before the onset of both tests, mice were deprived of any nesting material for 24 h. For nestlet shredding, mice were supplied with a nestlet (Ancare, Nassau, NY, USA) in their home cage for 60 min. Nestlets were weighed before and after the test, and the weight difference was used for calculations.

For nest building, mice were provided with a nestlet for 72 h and the nests were pictured after 20 and 48 h. Nests were scored using a published 5 point grading system^24^ with 1 being poor and 5 being the best.

### Locomotor Activity Assay

Locomotor activity was measured using a SuperFlex open field system (Omnitech Electronics, Accuscan, Columbus, OH), and a proprietary Fusion v6.5 software was used to track beam breaks to determine the total distance traveled during 60 min testing sessions. An injection (i.p) of saline was administered prior to testing. Following the injection, mice were placed in the sound-attenuated chamber, where they freely explored the chamber for 60 minutes. Distance traveled and time spent in zones (surround, center and corners) was calculated.

### Data Analysis

Statistical analyses were performed using Prism 10 (GraphPad Software Inc., La Jolla, CA). Data are reported as mean ± SEM. Data were analyzed using repeated measures analysis of variance (ANOVA) with group (water, fentanyl) and treatment (saline, naloxone) as between-subjects factors and time as the within-subjects factor, or two-way Student’s t-tests, where appropriate. Significant interaction effects were followed up using Bonferroni’s or Tukey’s post hoc tests.

## Results

To model oral fentanyl use in female C57BL/6J mice, we used a modified fDiD approach (Fig. 1A), in which mice were given 4-hour access to a single bottle of fentanyl (30 ug/ml in tap water) in their home cage 5 days a week for 5 weeks. Though fentanyl drinking mice drank less volume than their water control counterparts (Fig. 1B, 2-way RM ANOVA; main effect of group (F(1,38)=95.44), session (F(11.82,449.3)=2.828, P=0.0010) and group x session interaction (F(24,912)=1.742, P=0.0152)), there was a moderate increase in cumulative weekly fentanyl intake across the 5-week drinking paradigm (Fig. 1C, 1-way RM ANOVA, effect of week: F_(3.123, 59.33)_=5.573, P = 0.0017). Post hoc testing revealed significantly greater intake during week 5 compared with weeks 2 and 4, and a trend toward significance for week 3 compared with week 5 (P=0.0883). Furthermore, following the end of the 5^th^ session of week 4, fentanyl mice injected with naloxone displayed an increase in global somatic withdrawal score (including: fecal boli production, escape jumps, paw tremors, wet dog shakes, abnormal posture, grooming, and jaw tremors) compared to saline-injected fentanyl mice as well as saline- and naloxone injected water mice (Fig.1D). A 2-way ANOVA revealed significant effects of withdrawal treatment (F_(1, 36)_=212.9, P<0.0001) and fDiD solution (F_(1, 36)_=226.2, P, P<0.0001), as well as their interaction (F_(1, 36)_=159.4, P<0.0001). Tukey’s multiple comparisons showed a significant effect between Saline:Water, Naloxone:Water, and Saline:Fentanyl compared to the Naloxone:Fentanyl group (all 3 comparisons: P<0.0001). Collectively, these results suggest that the modified single-bottle fDiD paradigm supports sustained oral fentanyl drinking and produces physiological effects in female mice.

**Figure 1.**
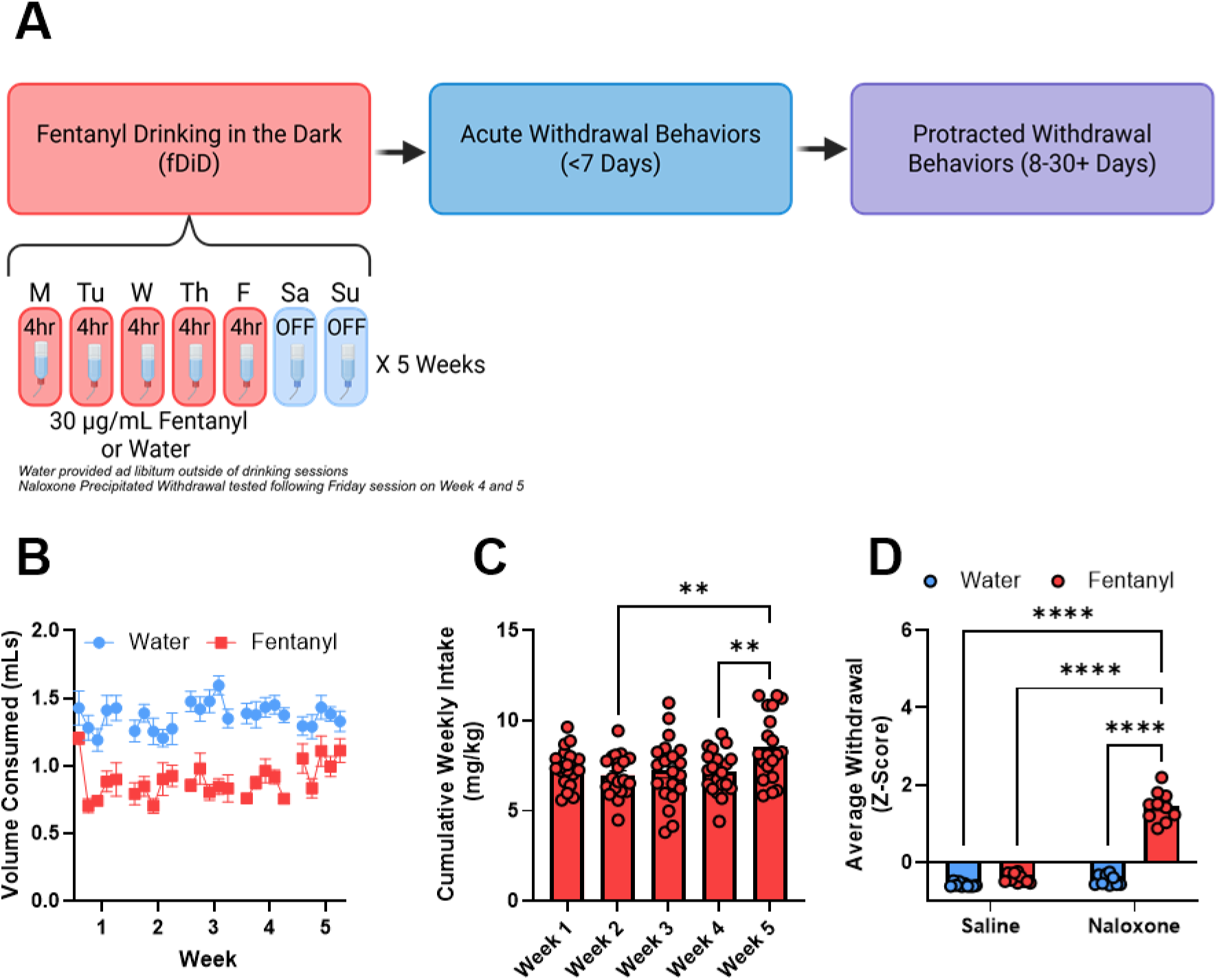
Chronic oral fentanyl consumption is sufficient to induce somatic withdrawal symptoms. (A) Experimental timeline of fentanyl DiD across 5 weeks. (B) Total fluid (fentanyl or water alone) consumed by fentanyl and water mice across the 5 weeks of DiD. (C) Cumulative weekly intake of fentanyl (mg/kg). Fentanyl mice consumed more fentanyl during week 5 compared to weeks 2 or 4. (D) Average withdrawal score following during the naloxone precipitated withdrawal showed fentanyl drinking mice that received naloxone exhibited more withdrawal behaviors than their counterparts. **P<0.1, ****P<0.0001

To determine whether chronic fentanyl exposure alters nociceptive sensitivity during drug exposure and abstinence, thermal and mechanical nociceptive behaviors were assessed longitudinally. We assessed thermal nociception and thermal hyperalgesia using the Hargreaves assay, which measures reflexive paw withdrawal latency to a radiant heat stimulus. Prior to the initiation of fentanyl drinking, we assessed baseline thermal sensitivity and found that paw withdrawal latencies did not differ between water- and fentanyl-drinking mice (Fig. 2B; unpaired t-test; t_38_=1.502, P=0.1414).

**Figure 2.**
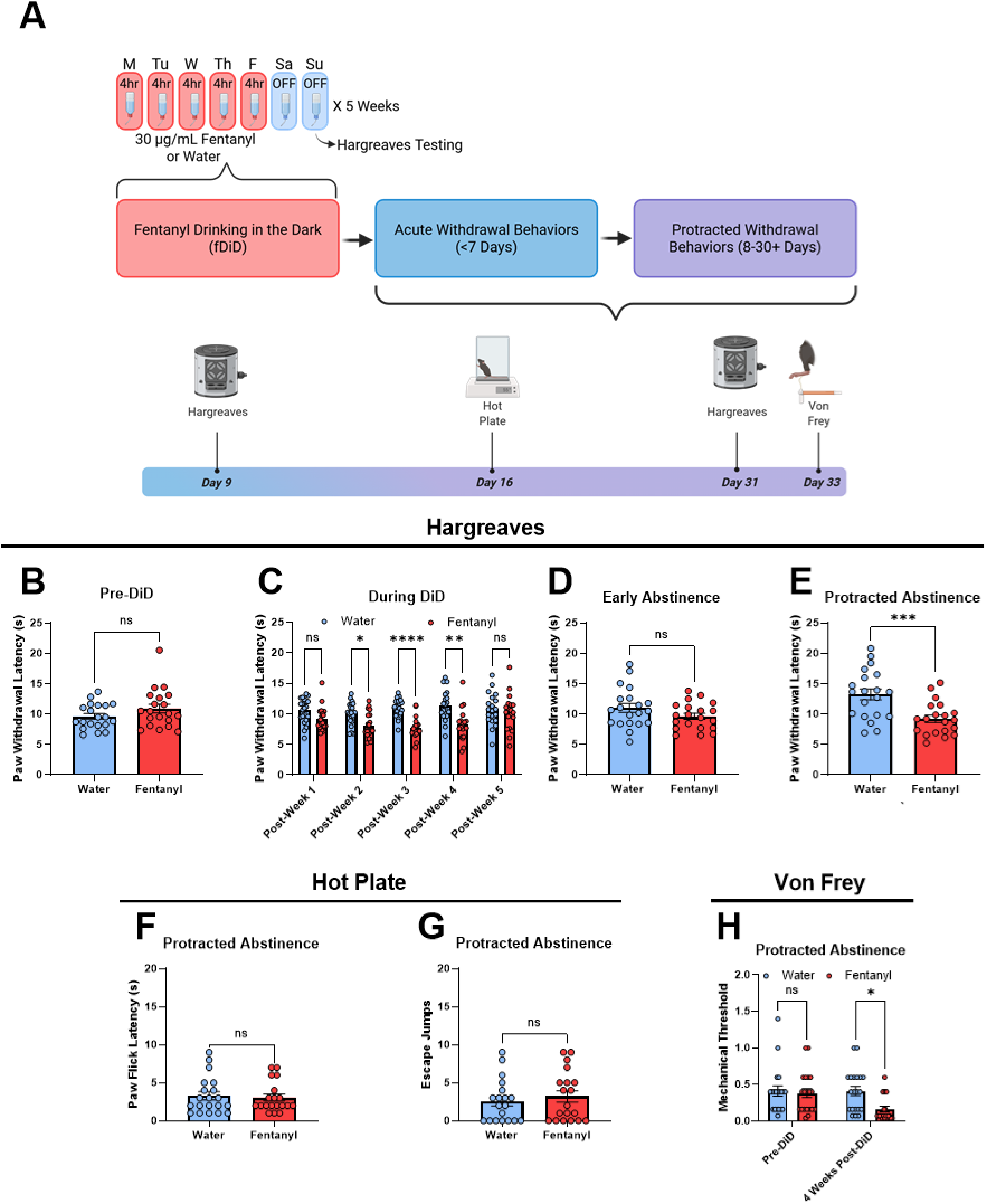
Pain sensitivity during fentanyl exposure, acute and prolonged withdrawal. (A) Experimental timeline of pain and nociceptive testing during fentanyl DiD and forced abstinence. (B) Prior to initiation of fDiD water and fentanyl mice displayed no difference in paw withdrawal latency on the Hargreaves assay. (C) During fDiD, mice 2 days following the final DiD session of each week. Fentanyl exposed mice displayed reduced paw withdrawal latencies following week 2, 3, and 4 of fentanyl drinking. (D) When tested a week following their final DiD session, no difference in paw withdrawal latency was observed. (E) However, when tested 30 days following the last session, fentanyl mice displayed reduced paw withdrawal latencies. (F) 2 weeks into forced abstinence, we tested response to a noxious thermal stimulus using the hot plate and found no difference in paw flick latency nor escape jumps (G). To assess mechanical sensitivity, we used the von Frey assay. Prior to DiD, there were no differences between water and fentanyl mice. (H) However, 30 days following abstinence there was a significant reduction in mechanical threshold in the fentanyl mice. *P<0.05, **P<0.01, ***P<0.001 ****P<0.0001

Thermal sensitivity was subsequently assessed following each week of the 5-week drinking paradigm. Repeated weeks of fentanyl drinking progressively altered thermal sensitivity, with fentanyl-drinking mice exhibiting significantly reduced paw withdrawal latencies following weeks 2, 3, and 4 compared with water controls (2-way RM ANOVA, treatment × week interaction: F_(3.531, 132.4)_=3.582, P=0.011; Bonferroni post hoc, Water < control, Week 2 P=0.0155, Week 3 P<0.001, and Week 4 P=0.0018; Fig. 2C), consistent with the development of thermal hyperalgesia during chronic fentanyl exposure. Interestingly, this reduction in withdrawal latency was no longer evident following the fifth week of drinking nor 9 days into abstinence (Fig. 2C and 2D). Notably, when tested after 30 days of abstinence, fentanyl-drinking mice again displayed significantly shorter withdrawal latencies than water controls fentanyl (Fig 2E; unpaired t-test; t_38_=3.675, P=0.0007), demonstrating the re-emergence of thermal hyperalgesia during protracted withdrawal.

We also accessed response to a noxious thermal stimulus using the hot plate assay 2 weeks into abstinence. When examining the response to the hot plate assay we found fentanyl-drinking mice did not differ from water controls in either paw flick latency (Fig. 2F) or the number of escape jumps (Fig. 2G), suggesting that thermal nociception has a temporally distinct trajectory across abstinence. addition, we tracked mechanical sensitivity longitudinally using the von Frey assay prior to initiation of drinking and 30 days into abstinence. Mechanical sensitivity did not differ between groups at baseline prior to fentanyl drinking; however, 1 month into abstinence, fentanyl drinking mice displayed a lower mechanical threshold compared to water controls (Fig. 2H, 2-way RM ANOVA, main effect of group F_(1, 38)_=4.976, P=0.0317, Bonferroni post hoc test: 4 weeks P=0.0101).

To determine whether acute withdrawal following chronic fentanyl exposure alters affective state or anxiety-like behavior, mice were evaluated within the first week following the final drinking session using the open field test, sucrose preference assay, and the elevated plus maze (Fig. 3A).

**Figure 3.**
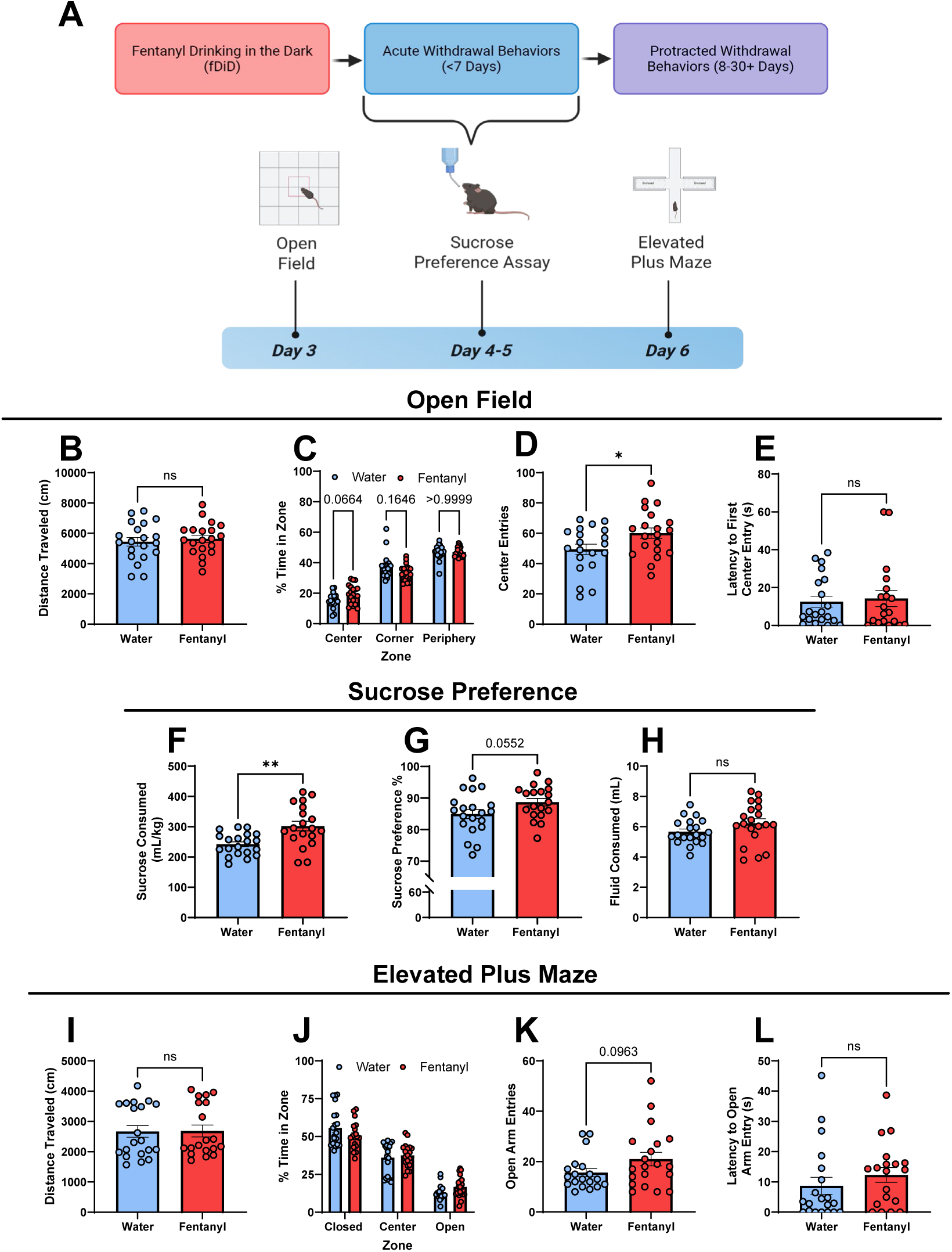
Chronic fentanyl drinking leads to emergence of affective and anxiety-like behavioral disturbances during early fentanyl abstinence. (A) Experimental timeline of affective and anxiety-like behavioral testing during early withdrawal (<7 days) of fDiD. On abstinence day 2, mice were test on the open field assay. Fentanyl drinking mice no difference in total distance traveled (B); however, the distribution of time spent in the zone of the maze were altered (C). Further, fentanyl mice showed greater center zone entries (D) but no difference in latency to enter the center (E). During the sucrose preference test (abstinence day 4-5), fentanyl exposed mice consumed more sucrose (F) and had a trend towards increased preference for sucrose over water (G) but no difference in total fluid consumed (H). On abstinence day XX, mice were test on the open field assay. Fentanyl drinking mice no difference in total distance traveled (E); however, the distribution of time spent in the zone of the maze were altered (F). Further, fentanyl mice showed greater center zone entries (G) but no difference in latency to enter the center (H). Lastly, mice were test on the EPM on abstinence day 6. There was no effect of drinking on total distance traveled (I) nor the distribution of time in the zones (J). However, there was a trend for increased open arm entries (K) and in fentanyl mice though no effect on latency to enter (L). *P<0.05, **P<0.01, ***P<0.001 ****P<0.0001

First, we assessed exploration of a novel environment using the open field assay. Total distance traveled did not differ between groups (Fig. 3B), indicating that chronic fentanyl exposure did not alter general locomotor activity during acute withdrawal. During the test, fentanyl mice displayed an altered pattern in the proportion of time spent across arena zones (Fig. 3C; 2-way RM ANOVA, main effect of zone: F_(1.538, 58.45)_ =201.6, P<0.0001, interaction of zone and group: F_(1.538, 58.45)_=3.881, P=0.0363). Bonferroni’s multiple comparisons test indicated a trend for fentanyl mice to spend more time in the center than water mice (P = 0.0664). Furthermore, fentanyl-drinking mice exhibited significantly more center entries than water controls (Fig. 3D; unpaired t-test, t_38_=2.205, P=0.0336), while center latency was unchanged (Fig. 3E). Together, these findings suggest that acute withdrawal was associated with increased exploration of the anxiogenic center of the arena without evidence of general locomotor changes, consistent with either reduced anxiety-like behavior or increased exploratory drive during early abstinence.

To evaluate whether acute withdrawal altered sensitivity to natural reward, mice were tested in a 24-h two-bottle sucrose preference assay. Fentanyl-drinking mice consumed significantly more sucrose than water controls (unpaired *t*-test, t_37_=3.437, P=0.0015; Fig. 3F), accompanied by a trend toward increased sucrose preference (Fig. 3G; unpaired *t*-test, P=0.0552), while total fluid consumption did not differ between groups (Fig. H). These findings indicate that acute withdrawal increased consumption of natural rewards and may reflect heightened reward seeking at early timepoints.

We then evaluated anxiety-like behavior using the EPM, an approach-avoidance conflict task^26^. Again, we found total distance traveled during the assay did not differ between groups, suggesting comparable locomotor activity (Fig. 3I). Likewise, fentanyl exposure did not alter the percentage of time spent in the closed, center, or open arms (Fig. 3J), and no differences were observed in either open-arm entries (Fig. 3H) or latency to enter the open arms (Fig. 3L). However, we did find a trend toward increased open-arm entries in fentanyl-drinking mice (Fig 3K; unpaired t-test, t_37_=1.706, P=0.0963).

Collectively, these findings indicate that female mice exhibit increased sucrose reward seeking and increases in exploratory behavior in anxiogenic environments.

Given the increased reward seeking and exploratory behaviors during acute withdrawal, we next asked whether affective and anxiety-like phenotypes emerge or evolve across prolonged abstinence.

Mice were tested in the acoustic startle assay 11 days after the final drinking session to assess startle reflex. The acoustic startle response measures the magnitude of reflexive responses to sudden auditory stimuli and is frequently used as an index of heightened arousal and stress responsiveness^27^. We found no group differences in the maximum startle response (Fig. 4B) or the latency to reach peak response across trials (Fig. 4C), suggesting no change in startle responsiveness or sensorimotor reactivity.

**Figure 4.**
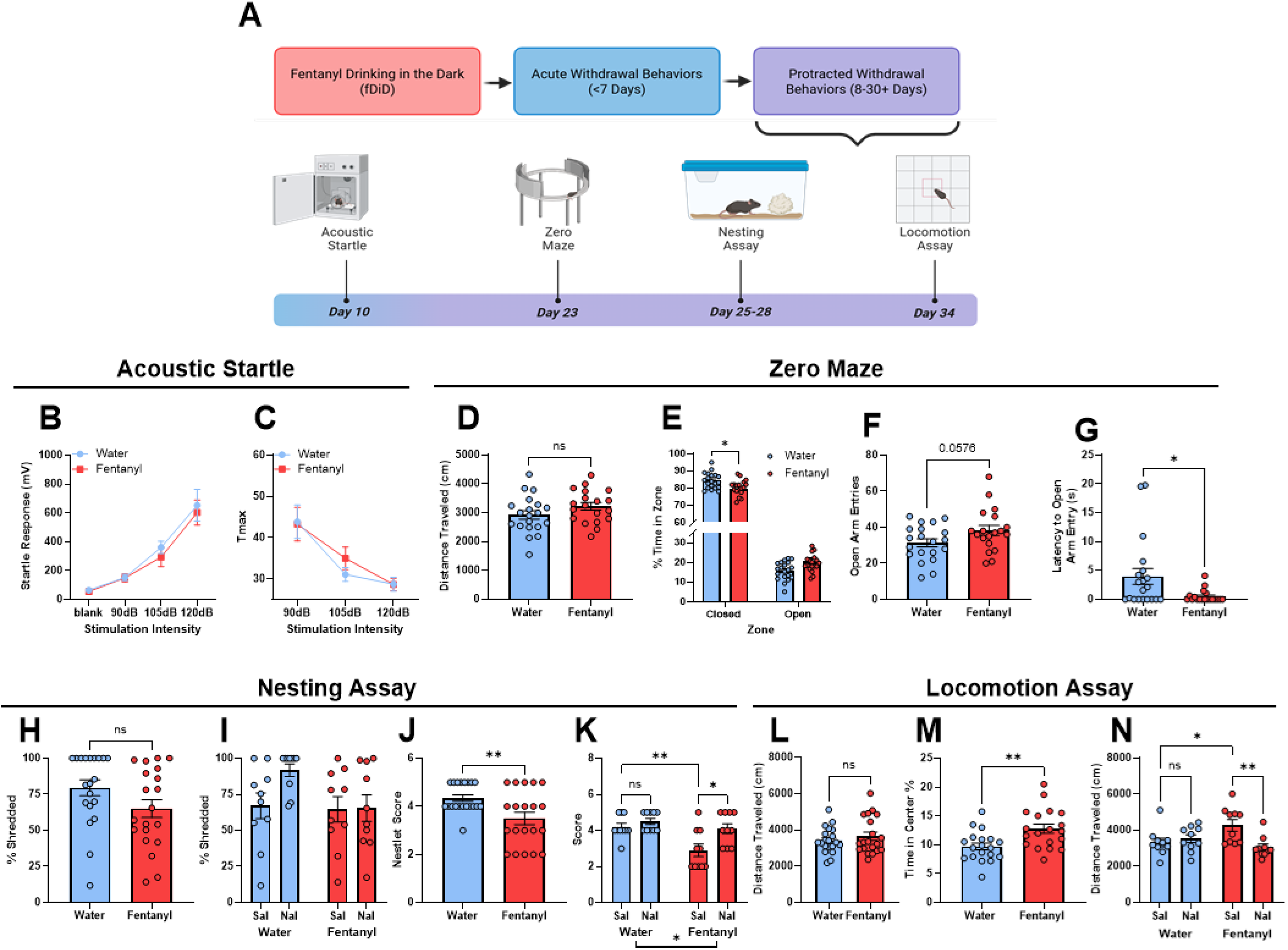
Chronic fentanyl drinking leads to emergence of affective and anxiety-like behavioral disturbances during early fentanyl abstinence. (A) Experimental timeline of affective and anxiety-like behavioral testing during protracted fentanyl abstinence (>7 days) of fDiD. Mice were tested on startle responsivity on abstinence day 11 using the acoustic startle assay. No differences were found in the maximum startle response (B) nor time to maximum startle response (C) between fentanyl or water consuming mice. On abstinence day 24, mice were tested on the zero maze. Fentanyl drinking mice no difference in total distance traveled (D); however, the distribution of time spent in the zone of the maze were altered (E), with fentanyl mice spending less time in the closed portion and more time in the open compartments. Further, fentanyl mice showed a trend towards greater open arm entries (F) and reduced latency to enter the open arm (G). Assessment of nesting behaviors was evaluated on abstinence day 26. We found no difference in the amount of nesting material shredded between drinking groups (H) and prior naloxone experience had no impact as well (I). However, fentanyl drinking mice had lower nest scores (J). When evaluating prior precipitated withdrawal experience, lower nesting scores were seen specifically mice that received saline compared to their water consuming counterparts (K). Further, fentanyl consuming mice that received naloxone displayed increased scores compared fentanyl-saline mice (K).. Lastly, mice were tested on locomotor activity on abstinence day 34. While no difference in total distance traveled was found between groups (L), fentanyl mice spent a greater percentage of time in the center portion of the arena (M). In evaluation of prior precipitated withdrawal experience, we found that in the first 10 minutes of the assay fentanyl drinking mice that received saline had greater locomotion than Water:Saline mice; however this hyperlomotion was not observed in naloxone receiving fentanyl mice (N). *P<0.05, **P<0.01

To evaluate anxiety-like behavior at a later abstinence timepoint, we tested mice on the elevated zero maze 24 days after the final drinking session. Of note, unlike the OFT and EPM, the elevated zero maze provides a continuous assessment of approach-avoidance behavior^28,29^ by eliminating the ambiguous center zone. As in prior assays, we found no difference in total distance traveled between groups (Fig. 4D). However, fentanyl-drinking mice spent significantly less time in the closed quadrants and more time in the open quadrants than water controls (Fig. 4E; 2-way RM ANOVA main effect of zone: F_(1, 37)_=1238; P<0.0001; interaction of zone and group: F_(1, 37)_=7.728, P=0.0085). A Bonferroni’s multiple comparisons test revealed reduced closed arm time (P=0.0138) and increased open arm time (P=0.0138) in fentanyl drinking mice compared to water. Fentanyl-exposed mice also exhibited a trend toward increased open-arm entries (Fig. 4F; unpaired t-test: t_37_=1.960, P=0.0576) and reduced latency to enter the open arm (Fig. 4G; unpaired t-test: t_37_=2.447, P=0.0193). Together, these findings indicate that the increased exploration of anxiogenic environments is maintained or potentially strengthened during prolonged abstinence when compared to the acute withdrawal period.

We next assessed nest-building behavior 26 days after the final drinking session, as nest-building is an ethologically relevant behavior that integrates motivational, cognitive, and motor functions and is disrupted in numerous neuropsychiatric and neurological disease models^30^. Following 24hr access to nesting material, fentanyl mice display a trend towards reduced shredding of the nestlet (Fig. 4H; unpaired t-test, t_38_=1.721, P=0.0933) and a significant reduction in nestlet composite score (Fig. 4I; unpaired t-test, t_38_=2.950, P=0.0054). Interestingly, unlike prior assays, when we evaluated the impact of prior naloxone exposure on the nestlet score, a 2-way ANOVA revealed a significant main effect of group (F_(1, 36)_= 10.88, P=0.0022), withdrawal drug treatment (F_(1, 36)_=8.473, P=0.0061), and a trend towards an interaction between the two factors (F_(1, 36)_=3.050, P=0.0893, Fig. 4J). A Tukey’s post hoc test showed a significant reduction in nest composite score in fentanyl-administered saline mice compared to water mice (P=0.0055), as well as an increase in nest score in naloxone-treated fentanyl mice compared to saline (P=0.0114). However, neither naloxone administration nor group had a significant impact on the percent of nestlet shredded during the assay (Fig. 4K). These data suggest that chronic fentanyl exposure produces deficits in species-typical motivated behaviors in protracted withdrawal that are sensitivity to withdrawal modality (precipitated vs spontaneous).

Finally, to examine whether the effects of nest building could result from increased exploratory locomotor activity during later, prolonged abstinence periods, we again tested mice for locomotor activity in a second novel environment approximately 1 month after the final drinking session. When comparing only the drinking-effect group, we found no difference in total distance traveled (Fig. 4L); however, fentanyl-treated mice displayed an increase in time spent in the center compared to water controls (Fig. 4M; unpaired t-test, t_36_=3.110, P=0.0037). However, again, when we evaluated the impact of prior withdrawal drug exposure, a 2-way RM ANOVA of the first 10 minutes of the assay revealed a significant effect of withdrawal treatment drug (Fig. 4N, F_(1, 36)_=4.352, P=0.0441) and interaction of group and withdrawal treatment (F_(1, 36)_= 8.587, P=0.0058). A Tukey’s multiple comparisons test showed a significant increase in locomotion in saline-treated fentanyl mice compared to saline-treated water mice (P=0.0457), and a reduction in locomotion in naloxone-treated fentanyl mice compared to saline-treated fentanyl mice (P=0.0058). Together, these findings demonstrate that locomotor activity during prolonged abstinence is influenced by both chronic fentanyl exposure and prior naloxone-precipitated withdrawal, supporting the notion that withdrawal history may shape behavioral adaptations well beyond the acute withdrawal period.

## Discussion

The present study employed a modified home-cage oral fentanyl drinking paradigm to characterize behavioral adaptations associated with chronic fentanyl exposure and withdrawal in female mice. Consistent with prior reports using DiD fentanyl drinking paradigms, female mice exhibited robust naloxone-precipitated somatic withdrawal following the final fDiD session, confirming they consumed dependency-producing levels of fentanyl. We expanded on these studies and demonstrate that the behavioral consequences of chronic fentanyl exposure are jointly shaped by three interacting variables: history of fentanyl exposure, precipitated withdrawal experience, and duration of abstinence. Collectively, these results support a model of withdrawal that emerges as a dynamic process in which nociceptive, affective, and motivated behaviors evolve across distinct phases of drug exposure and abstinence. Further, these findings provide one of the first longitudinal behavioral characterizations of chronic oral fentanyl exposure in female mice and highlight the importance of considering both withdrawal and abstinence duration when modeling OUD.

Using a modified fDiD approach, we found that female mice consume steady volumes of fentanyl across weeks with a mild increase at week 5. However, the volume of fentanyl consumed was consistently lower than that of water by the control mice. A prior study found that when offered increasing concentrations of fentanyl, volume consumed decreased, suggesting that mice may titrate the volume consumed^13^. Another study found that mice consumed more fentanyl in an operant-based format when compared to fDiD homecage drinking and suggested this “binge” drinking model^31^ may drive high consumption at a rapid pace that may restrict total volume consumed^14^. Moreover, we found that administration of naloxone following 4 weeks of fDiD led to robust precipitated withdrawal symptoms, demonstrating oral fentanyl drinking was sufficient to attain physiologically relevant levels of fentanyl intake.

Hyperalgesia and reduced pain tolerance are both commonly cited outcomes of opioid withdrawal^32–34^ and suggested to drive increased relapse propensity^15^. Women report higher rates of chronic pain and are more likely to receive prescription opioids, factors that increase exposure to opioid medications and contribute to risk of misuse^35^. Moreover, recent work demonstrated that biological sex and ovarian hormones modulate the interaction between pain and fentanyl consumption, underscoring the importance of studying pain-related outcomes during fentanyl abstinence in females^36^. An interesting finding of the present study was that pain-related behavioral adaptations followed a distinct temporal trajectory across fentanyl exposure into acute and protracted withdrawal. Thermal hyperalgesia emerged during weeks two through four of fentanyl drinking, normalized during the final week of exposure despite continued fentanyl access, disappeared during early abstinence timepoints, and re-emerged during prolonged abstinence. Further, mechanical nociceptive sensitivity was also heightened after over a month of forced abstinence. Together, these findings suggest that different nociceptive modalities are affected by chronic fDiD and abstinence across unique timescales following chronic fentanyl exposure.

During early abstinence (<7 days) we found that fentanyl drinking mice displayed increased sucrose consumption and increased exploratory behaviors in anxiogenic environments on the EPM and OFT. Our observations during acute withdrawal also differ somewhat from several experimenter-administered opioid paradigms, in which robust anxiety-like behaviors and anhedonia often emerge within the first 3-5 days following drug cessation^37–40^ that can last for weeks into abstinence^20,41–44^. While methodological differences, including route of administration, dosing strategy, pattern of opioid exposure, and sex, likely contribute to these discrepancies, they collectively suggest that withdrawal-associated behaviors may be context-dependent. Mirroring a previous study using an identical oral fentanyl drinking model^13^, we also observed increased sucrose preference in female mice, suggesting that chronic oral fentanyl drinking may engage neuroadaptive mechanisms distinct from those induced by repeated passive opioid administration. In addition, a consistent behavioral finding throughout abstinence was increased exploration of anxiogenic environments. During acute withdrawal, fentanyl-exposed mice exhibited increased center entries in the open field and modest trends toward greater open-arm exploration in the EPM. These effects became more pronounced during prolonged abstinence, with females spending more time in the open quadrants of the EZM. Although these behaviors are traditionally interpreted as reduced anxiety-like behavior, these findings may reflect behavioral disinhibition, altered exploratory behavior, or changes in decision-making processes following chronic fentanyl exposure. This notion is supported by evidence demonstrating that oral fentanyl exposure selectively impairs long-term fear extinction memory encoding or retrieval following fear conditioning^13^ and separate work demonstrating chronic oral fentanyl reinforcement increases perseverative responding and promotes a shift toward habitual responding for both drug and nondrug rewards, particularly in females^45^.

Interestingly, only the nesting behaviors captured nearly 4 weeks following the last fentanyl session had a significant effect of prior naloxone treatment. Fentanyl consuming mice that experience multiple naloxone-precipitated withdrawals had better nesting scores compared to saline-treated fentanyl drinking mice. When evaluating if changes in nesting behavior coincide with changes in general locomotion, we also found that naloxone administration in fentanyl drinking mice reduced hyperactivity seen in saline treated fentanyl mice. Together, these findings suggest that repeated naloxone-precipitated withdrawal may differentially shape the long-term behavioral consequences of fentanyl abstinence, potentially engaging neuroadaptive processes that selectively influence pain sensitivity, ethologically relevant behaviors, and locomotor activity.

Prior work from our group demonstrated that ventrolateral periaqueductal gray (vlPAG)/dorsal raphe (DR) dopamine circuits regulate pain-related behaviors in a sex-dependent manner, with males preferentially exhibiting anti-nociceptive responses while females display alterations in locomotor responses to salient stimuli rather than equivalent reductions in nociceptive sensitivity^22^. Combined with additional evidence of heightened glutamatergic drive onto dorsal raphe dopamine neurons during protracted morphine withdrawal^46^, these findings raise the possibility that persistent adaptations within vlPAG/dorsal raphe dopamine circuits represent a convergent mechanism underlying the time- and context-dependent behavioral responses observed during protracted fentanyl abstinence. Although the present study did not directly investigate the neural circuitry underlying fentanyl withdrawal, the evolving nociceptive phenotype observed across fentanyl drinking and abstinence is consistent with the broader concept that chronic opioid exposure reshapes adaptive behavioral responses to aversive stimuli rather than producing a static pain-like phenotype. Rather than broadly increasing nociceptive sensitivity, chronic fentanyl exposure may alter how females integrate nociceptive information with motivational and behavioral outputs.

In conclusion, our findings demonstrate that voluntary oral fentanyl exposure in female mice produces dynamic behavioral adaptations that extend well beyond the period of drug exposure and evolve across distinct phases of abstinence. Rather than exhibiting a uniform withdrawal phenotype, females displayed temporally dissociable changes in nociception, exploratory behavior, and ethologically relevant behaviors, indicating that chronic fentanyl exposure may engage diverse neuroadaptive processes. Furthermore, these observations underscore the importance of considering both the duration of abstinence and prior withdrawal experience when modeling OUD, as behavioral outcomes differed substantially across time despite identical drug exposure histories. Ultimately, these findings provide an important framework for investigating the neural mechanisms underlying protracted opioid abstinence and may facilitate the development of sex-informed therapeutic strategies aimed at alleviating persistent withdrawal symptoms and reducing relapse vulnerability.

## Notes

### Competing Interest Statement

The authors have declared no competing interest.

